# Using plant traits to understand the contribution of biodiversity effects to community productivity in an agricultural system

**DOI:** 10.1101/2020.06.17.157008

**Authors:** Nadine Engbersen, Laura Stefan, Rob W. Brooker, Christian Schöb

## Abstract

- Increasing biodiversity generally enhances productivity through selection and complementarity effects not only in natural but also in agricultural systems. However, explaining why diversity enhances productivity remains a central goal in agricultural science.
- In a field experiment, we constructed monocultures, 2- and 4-species mixtures from eight crop species with and without fertilizer and both in temperate Switzerland and semi-arid Spain. We measured environmental factors and plant traits and related these in structural equation models to selection and complementarity effects to explain yield differences between monocultures and mixtures.
- Increased crop diversity increased yield in Switzerland. This positive biodiversity effect was driven to almost same extents by selection and complementarity effects, which increased with plant height and C:N ratio, respectively. Also, ecological processes driving yield increases from monocultures to mixtures differed from those responsible for yield increases through the diversification of mixtures.
- While selection effects were mainly driven by one species, complementarity effects were linked to higher productivity per unit N. Yield increases due to mixture diversification were driven only by complementarity and were not mediated through the measured traits, suggesting that ecological processes beyond those measured in this study were responsible for positive diversity effects on yield beyond 2-species mixtures.

## 1. Introduction

Plant primary productivity increases with higher species diversity (e.g. Tilman et al. (2001); Cardinale et al. (2006)). While the majority of research in this area has been done in perennial systems (Tilman et al. 2001; Weisser et al. 2017; Huang et al. 2018), recent studies have demonstrated similar effects in annual systems (Li et al. 2014; Brooker et al. 2015; Stomph et al. 2020). Intercropping of annual crops, where at least two crop species are grown in close proximity at the same time, is therefore a promising application of agroecological concepts. Intercropping can make use of species complementarity and beneficial above- and belowground facilitation and niche differentiation, which can ultimately lead to overyielding, i.e. increased yield in mixture compared with the average of the monocultures (Vandermeer 1989).

Two main mechanisms have been proposed to explain positive biodiversity–productivity relationships. First, sampling or selection effects (hereafter: selection effects) encompass the greater probability that more diverse communities include highly productive species or functional groups, which then account for the majority of productivity (Huston 1997; Tilman et al. 1997). Enhanced ecosystem functioning in diverse agroecosystems can be driven by selection effects (i.e. communities with more species are more likely to host a high-performing species). For instance, in China, a hotspot of intercropping, a recent meta-analysis has shown that 10% of all yield gain of intercropping compared to sole cropping was due to selection effects (Li et al. 2020).

The second mechanism is complementarity through resource partitioning or facilitation. Resource partitioning involves more diverse communities containing species with contrasting demands on resources, which leads to a more complete exploitation of available resources in diverse plant communities compared with monocultures and hence increased productivity (Tilman et al. 1997; Loreau and Hector 2001). Facilitation involves plants altering their environment in a way that is beneficial to at least one co-occurring species (Brooker et al. 2008). However, knowledge about the precise mechanisms that lead to overyielding in crop mixtures and how environmental conditions can alter these mechanisms still remains incomplete (Duchene et al. 2017).

Resource partitioning can occur across spatial, temporal or chemical gradients. For example, different rooting depths allow plants to take up water or nutrients from different soil layers, thus limiting competition. While this has been observed for water uptake (Miyazawa et al. 2009), the evidence of resource partitioning for soil nutrients as a driver of biodiversity effects is less clear (Von Felten et al. 2012; Jesch et al. 2018). Aboveground, diverse communities have been observed to harbor more diverse canopy growth forms allowing for a more complete use of photosynthetically active radiation (PAR) (Spehn et al. 2002; Fridley 2003), leading in some cases to yield advantages in mixtures compared with monocultures (Bedoussac and Justes 2009).

Facilitation involves plants altering their environment in a way that is beneficial to at least one co-occurring species (Brooker et al. 2008). Facilitation among plants can happen via either the enrichment of resource pools by one plant for neighboring plants or the mediation of physical stress.The facilitative benefits of legume neighbors for enhancing soil N are well known, for example for cereals intercropped with legumes (Jensen 1996; Hauggaard-Nielsen et al. 2001; Spehn et al. 2002; Temperton et al. 2007). Other below-ground facilitative mechanisms shown to occur in intercrops include enrichment of resource pools by hydraulic lift, which not only facilitates water uptake (Sekiya and Yano 2004), but can also enhance nutrient mobilization and lead to increased nutrient status of the intercrop (Sun et al. 2013). Other evidence of facilitation mediating physical stress in crop systems come from studies of barley variety mixtures, where a denser canopy structure, shading of the soil surface, and thus reduced evaporation were observed to decrease the soil temperature due (Cooper et al. 1987). Also, plant species can alleviate the microclimate for their neighbors by mediating wind, heat or photoinhibition (Wright et al. 2017).

However, despite these examples of different resource partitioning and facilitation mechanisms occurring in crop systems, knowledge about the precise mechanisms that lead to overyielding in crop mixtures, and how environmental conditions can alter these mechanisms, still remains incomplete (Duchene et al. 2017). While there is abundant evidence for the presence of facilitation and complementarity in diverse agricultural systems, the presence of these processes alone does not guarantee overyielding.

Here, we applied ecological knowledge and methods to agriculture by assessing to what extent complementarity and selection effects drive yield gains in crop mixtures compared to crop monocultures and how these effects are related to plant functional traits and environmental factors. By linking frequently-used plant traits to yield gains in mixtures, we can improve our ability to predict optimal species combinations which can help to promote sustainable agricultural production through intercropping. We used four plant traits indicative of resource use: leaf dry matter content (LDMC), specific leaf area (SLA), carbon to nitrogen ratio (C:N ratio) and plant height and two environmental factors: PAR and volumetric soil water content (VWC). We used a structural equation model (SEM) to assess whether overyielding is driven by selection or complementarity effects or by a combination of both. Furthermore, we used this hierarchical model to understand the context–dependence of the complementarity and selection effects and how they are linked to plant functional traits.

## 2. Materials and Methods

### 2.1. Site description

The study was carried out in two outdoor experimental gardens in Zürich, Switzerland and Torrejón el Rubio, Cáceres, Spain. The Swiss site was located at an altitude of 508 m a.s.l. (47°23’45.3” N 8°33’03.6” E), the Spanish site at 290 m a.s.l. (39°48’47.9” N 6°00’00.9” W). The regional climate in Switzerland was classified according to Köppen-Geiger (Kottek et al. 2006) as warm temperate, fully humid with warm summers while in Spain it was classified as warm temperate, summer dry with hot summers. Precipitation in Zürich during the growing season 2018 (April – August) was 572 mm, and in Spain 218 mm (February – June). Daily average hours of sunshine during the growing season were 5.8h in Zürich and 8.4h in Cáceres. Temperatures between the two sites were similar: average of daily mean, minimum and maximum temperatures were 14.0 °C, 9.3 °C and 18.6°C in Zürich and 14.6 °C, 9.6°C and 19.6°C in Cáceres. All climatic data are from the Deutsche Wetterdienst (www.dwd.de).

The experimental garden at each location covered 98 m^2^, divided into 392 square plots of 0.25 m^2^ and 40 cm depth. However, plots were open at the bottom and allowed root growth beyond 40 cm. In Switzerland, the plots were arranged in 14 beds of 7 × 1 m, with two rows of 14 plots, resulting in 28 plots per bed. In Spain, the plots were arranged in 14 beds of 10 × 1 m, with two rows of 20 plots, resulting in 40 plots per bed. The plots were filled with local, unenriched agricultural soil. Soil structure and composition therefore differed between the sites. In Switzerland, soil was composed of 45% sand, 45% silt, 10% clay and contained 0.19% nitrogen, 3.39% carbon, and 333 mg total P/kg with a mean pH of 7.25. Spanish soils consisted of 78% sand, 20% silt, 2% clay and contained 0.05% nitrogen, 0.5% carbon, 254 mg total P/kg with a mean pH of 6.3.

The experimental gardens were irrigated throughout the growing season with the aim of maintaining the differences in precipitation between the two sites but assuring survival of the crops during drought periods. In Switzerland, the dry threshold was set to 50% of field capacity, with a target of 90% of field capacity, while in Spain the thresholds were 17% and 25% of field capacity, respectively. Automated irrigation was configured such that irrigation would start if the dry threshold was reached, and irrigate until the target threshold was reached.

At each site, half of the beds were chosen randomly to be fertilized with N-P-K (1-1.7-1) while the other half served as unfertilized controls. Fertilizer was applied three times: 50 kg/ha just before sowing, another 50 kg/ha when wheat was tillering and 20 kg/ha when wheat was flowering.

### 2.2. Crop species and cultivars

At each site, experimental communities were constructed with eight annual crop species of agricultural interest. The crop species belonged to four different phylogenetic groups: *Triticum aestivum* (wheat, C3 grass) and *Avena sativa* (oat, C3 grass), *Lens culinaris* (lentil, legume) and *Lupinus angustifolius* (lupin, legume), *Linum usitatissimum* (flax, herb [superrosids]) and *Camelina sativa* (false flax, herb [superrosids]), *Chenopodium quinoa* (quinoa, herb [superasterids]) and *Coriandrum sativum* (coriander, herb [superasterids]). The four phylogenetic groups were based on their phylogenetic distances: Cereals diverged from the other groups 160 million years ago (mya); superasterid herbs diverged from superrosid herbs including legumes 117 mya and finally, legumes diverged from the other superrosid herbs 106 mya (*TimeTree*). Phylogenetic distance was chosen as a criterion for functional similarity as it is often positively correlated with functional diversity and acts as a proxy to assess the impacts of species diversity on ecosystem functions (Mouquet et al. 2015). At both sites, we grew commercial cultivars typically used for organic farming in Switzerland (Table S1). While these were bred for a Swiss climate, their cultivation in Spain demonstrated the ability of these cultivars to adapt to a climate change-type scenario, with conditions considerably warmer and drier than in Switzerland.

### 2.3. Experimental crop communities

Experimental crop communities at both sites consisted of eight different monocultures, 24 different 2-species mixtures consisting of two different phylogenetic groups and 16 different 4-species mixtures consisting of four different phylogenetic groups. Every combination of 2-species mixture with two species from different phylogenetic groups and every possible 4-species mixture with species from four different phylogenetic groups were planted. Each experimental community was replicated two times. Plots were randomized within each country and fertilizer treatment. Each monoculture and mixture community consisted of one, two or four crop species planted in four rows. The row order of the species was chosen randomly. Sowing densities differed among phylogenetic groups and were based on current cultivation practice (Table S1). Sowing was done by hand in early February 2018 in Spain and early April 2018 in Switzerland.

### 2.4. Data collection

Leaf traits were measured at the time of flowering (May 2018 in Spain, 94 days after sowing; June 2018 in Switzerland, 65 days after sowing). Three individuals per crop species per plot were randomly marked and their height measured with a ruler from the soil surface to the highest photosynthetically active tissue. Of each marked individual, 1 to 10 healthy leaves were sampled and immediately wrapped in moist cotton and stored overnight at room temperature in open plastic bags. For the subsequent leaf trait measurements (specific leaf area [SLA] and leaf dry matter content [LDMC]) we followed standard protocols (Cornelissen et al. 2003). The leaf samples were blotted dry to remove any surface water and immediately weighed to produce a value for saturated weight. They were instantly scanned with a flatbed scanner (CanoScan LiDE 120, Canon) and oven-dried in a paper envelope at 80 °C for at least 48 h. After drying, the leaf samples were reweighed to obtain dry weight. Values for dry matter content were calculated as the ratio of leaf dry mass (mg) to saturated leaf mass (g). The leaf scans were analyzed for leaf area with the image processing software ImageJ 1 (Rasband 1997-2018). SLA was calculated as the ratio of leaf area (cm^2^) to dry mass (g) and is a measure of the investment in dry mass per light-intercepting surface and used to assess light acquisition strategies; higher SLA values can be observed under low light conditions as a larger leaf area per leaf mass (= thinner leaves) enables increased surface for light capture at lower C costs (Weiher et al. 1999; Evans and Poorter 2001). LDMC is often inversely related to SLA but also includes soil-water relations and hence indicates impacts of resource conditions, including water (Pérez-Harguindeguy et al. 2013)

At the time of harvest (duration of crop growth from sowing to harvest given in table S2) all crops in each plot were harvested. The three marked individuals used for the trait measurements were collected separately, while all remaining plants per crop species per plot were pooled together. Plant shoots were cut at the soil surface and biomass and seeds were separated. The total number of individuals per crop species per plot was recorded. Fruits were air-dried. Afterwards, seeds were separated from chaff with a threshing machine (in Switzerland: Allesdrescher K35, Baumann Saatzuchtbedarf, Germany; in Spain: Hege 16, Wintersteiger, Austria). Vegetative biomass was oven-dried at 80 °C until constant weight.

Interception of PAR by the plant canopy was measured weekly with a LI-1500 (LI-COR Biosciences GmbH, Germany). In each plot, three PAR measurements were taken around noon by placing the sensor on the soil surface in the center of each of the three in-between rows. Light measurements beneath the canopy were put into context through simultaneous PAR measurements of a calibration sensor, which was mounted on a vertical post at 2 m above ground in the middle of the experimental garden. FPAR (%) indicates the percentage of PAR that was intercepted by the crop canopy. VWC in the soil was measured weekly with a ML3 ThetaProbe Soil Moisture Sensor (Delta-T, Cambridge). The measurements were taken in the center of each of the three in-between rows per plot. For further data analysis, we used FPAR and VWC values from the week of leaf trait measurements (Spain: 92 days after sowing; Switzerland: 62 days after sowing).

### 2.5. Plant N analyses

For chemical analyses of the plant tissue we pooled the three dried leaf samples of the marked individuals per plot and per crop species. Leaf samples were ground for 20 minutes in 1.2 ml tubes with two stainless steel beads in a bead mill (TissueLyserII, Qiagen). Afterwards, either 100 mg (if available in sufficient amount) or 4 mg (+/-1 mg) (if the sample was too small) of ground leaf material were weighed into tin foil cups or 5 × 9 mm tin capsules and analyzed for C and N contents. The 400 large samples were analyzed on a LECO CHN628C elemental analyzer (Leco Co., St. Joseph, USA) and the 505 small samples on a PDZ Europa 20-20 isotope ratio mass spectrometer linked to a PDZ Europa ANCA-GSL elemental analyzer (Sercon Ltd., Cheshire, UK), respectively. Eight samples were cross-referenced on both analytical devices (Fig. S1) and measured values from LECO were corrected to account for the differences between the devices (correction factors are 1.0957 for N and 1.026 for C, respectively).

### 2.6. Data analysis

Prior to data analysis the dataset was screened for missing data or incorrect data recording and the whole plot was removed when any of these occurred. A total of 467 plots remained, 231 in Switzerland and 236 in Spain.

All statistical analyses were performed in R version 3.6.0. (R Core Team 2019). We used general linear mixed-effects models using restricted maximum likelihood estimation to explain yield at the community-level. We assessed the significance of the fixed effects using type-I ANOVA and the Satterthwaite approximation of denominator degrees of freedom (lme4 (Bates et al. 2015) and lmerTest (Kuznetsova et al. 2017) packages). Yield was log transformed. The fixed effects of the model were country (Switzerland versus Spain), fertilizer (fertilized versus unfertilized), species number (2 versus 4) nested in diversity (monocultures vs mixtures), and interactions among the fixed effects (except between the nested terms). Random terms were species composition, garden bed and the interactions of garden bed with diversity and garden bed with species number. To test for effects of diversity within each country, we performed pairwise multiple comparison Tukey tests.

We used the same linear mixed-effects models as mentioned above to analyze treatment effects on environmental factors (FPAR, VWC), plant traits (SLA, LDMC, height, leaf N, C:N ratio) and biodiversity effects (selection and complementarity effects) with country, fertilizer, species number nested in diversity and interactions among these as fixed effects. Random terms were species composition, garden bed and their interaction. Response variables were log transformed, except for biodiversity effects.

Using structural equation modeling (SEM) we analyzed the sign and intensity of treatment effects and abiotic factors on plant traits, biodiversity effects and community-level yield, based on our complex *a priori* model (Fig. S2). To explain differences in community-level yield between mixtures and monocultures, we calculated Δyield as the difference between the summed community-level yields of all species in a mixture plot and the average of the mean community-level yields of all monocultures corresponding to the species in the mixture plot. Thus, Δyield compares the observed yield in the mixture with the expected yields in a mixture based on their yields in a monoculture. Δyield was square root transformed to meet the assumptions of normal distribution. We quantified the biodiversity effect (Δyield) and its two components, the complementarity and selection effects according to Loreau and Hector (Loreau and Hector 2001).

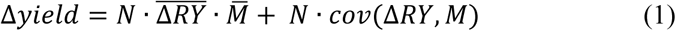

Where N is the number of species in the plot. ΔRY is the deviation from expected relative yield of the species in mixture in the respective plot, which is calculated as the ratio of observed relative yield of the species in mixture to the yield of the species in monoculture. The first component of the biodiversity effect equation 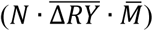 is the complementarity effect, while the second component (*N* · *cov*(Δ*RY, M*)) is the selection effect. We calculated ΔVWC and ΔFPAR according to equation 2.

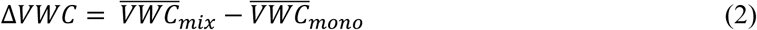

Where 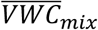 is the average of all three measurements of VWC per mixture plot and 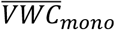 the average of all three measurements of VWC of the respective monoculture plots. The same was calculated for ΔFPAR. To scale up plant trait measurements to the community level, community-weighted means in mixtures and monocultures were used to calculate ΔSLA, ΔLDMC, ΔC:N ratio and Δheight:

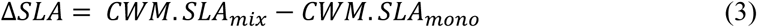

Where CWM.SLA_mix_ is the community-weighted mean of SLA of all species in a mixture plot and CWM.SLA_mono_ the community-weighted mean of SLA in monoculture of all the species in the respective plot. Aboveground biomass of each species was used as weights. ΔLDMC, ΔC:N ratio and Δheight were calculated likewise according to formula 3. The SEM was constructed using LAVAAN (Rosseel 2012) in R. For each variable, we report the conditional coefficient of determination (R^2^_c_), which represents the variation explained by fixed and random effects and was calculated with the MUMIN package (Bartoń 2019).

## 3. Results

### 3.1. Response of community-level yield to treatments

Community-level yield was significantly affected by country, with 88% higher yields in Switzerland compared with Spain (Fig. **1a**, Table 1). The addition of fertilizer did not show any effect on yield in either country nor on either diversity level. Mixtures produced significantly more yield than monocultures –and 4-species mixtures yielded more than 2-species mixtures (Fig. **1a**, Table 1). Tukey post hoc tests revealed that productivity of mixtures was enhanced, particularly in Switzerland (Table S3), where 4-species mixtures yielded 30% more than 2-species mixtures and 93% more than monocultures. Also, 2-species mixtures in Switzerland yielded 48% more than monocultures. In Spain, yield did not respond significantly to different diversity levels. The presence of a legume in mixtures increased yields in Spain by 72% but decreased it by 23% in Switzerland (Fig. **1b**, Table S3). The three-way interaction of country × fertilizer × legume indicates that mixtures without legumes yielded more in Switzerland in both fertilizer treatments, while in Spain mixtures with legumes had higher yields, but only in unfertilized soils (Fig. **1b**, Table S3).

**Table 1:** Results of mixed effects ANOVA testing effects of the different treatments (country, fertilizer, legume, diversity (monocultures vs. mixtures) and mixture diversification (2-vs. 4-species mixtures) on community-level yield. SS: Sum of squares, MS: mean of squares, numDF: degrees of freedom of term, denDF: degrees of freedom of error term, F-value: variance ratio, P: error probability. P-values in bold are significant at α = 0.05 (* P < 0.05, ** P < 0.01, *** P < 0.001). n = 315

|  | SS | MS | numDF | denDF | F-value | P |
| --- | --- | --- | --- | --- | --- | --- |
| **country** | 4.691 | 4.691 | 1 | 24.98 | 12.64 | **0.002\*\*** |
| *fertilizer* | 1.192 | 1.192 | 1 | 23.75 | 3.212 | 0.086 |
| *legume* | 0.201 | 0.201 | 1 | 43.24 | 0.542 | 0.466 |
| **diversity** | 1.524 | 1.524 | 1 | 42.67 | 4.107 | **0.049\*** |
| **mixture diversification** | 1.933 | 1.933 | 1 | 43.78 | 5.209 | **0.027\*** |
| *country × fertilizer* | 0.384 | 0.384 | 1 | 23.65 | 1.035 | 0.319 |
| **country × legume** | 16.99 | 16.99 | 1 | 257.1 | 45.77 | **8.91E-1\*\*\*** |
| **country × diversity** | 15.01 | 15.01 | 1 | 79.23 | 40.45 | **1.20E-08\*\*\*** |
| **country × mixture diversification** | 24.95 | 24.95 | 1 | 246.2 | 67.22 | **1.34E-14\*\*\*** |
| *fertilizer × legume* | 0.356 | 0.356 | 1 | 254.9 | 0.958 | 0.329 |
| *fertilizer × diversity* | 0.059 | 0.059 | 1 | 76.7 | 0.158 | 0.692 |
| *fertilizer × mixture diversification* | 0.014 | 0.014 | 1 | 244.7 | 0.037 | 0.849 |
| **country × fertilizer × legume** | 1.518 | 1.518 | 1 | 253.9 | 4.089 | **0.044\*** |
| *country × fertilizer × diversity* | 0.488 | 0.488 | 1 | 77.59 | 1.315 | 0.255 |
| *country × fertilizer × mixture diversification* | 0.997 | 0.997 | 1 | 245.5 | 2.685 | 0.103 |

**Figure 1:**
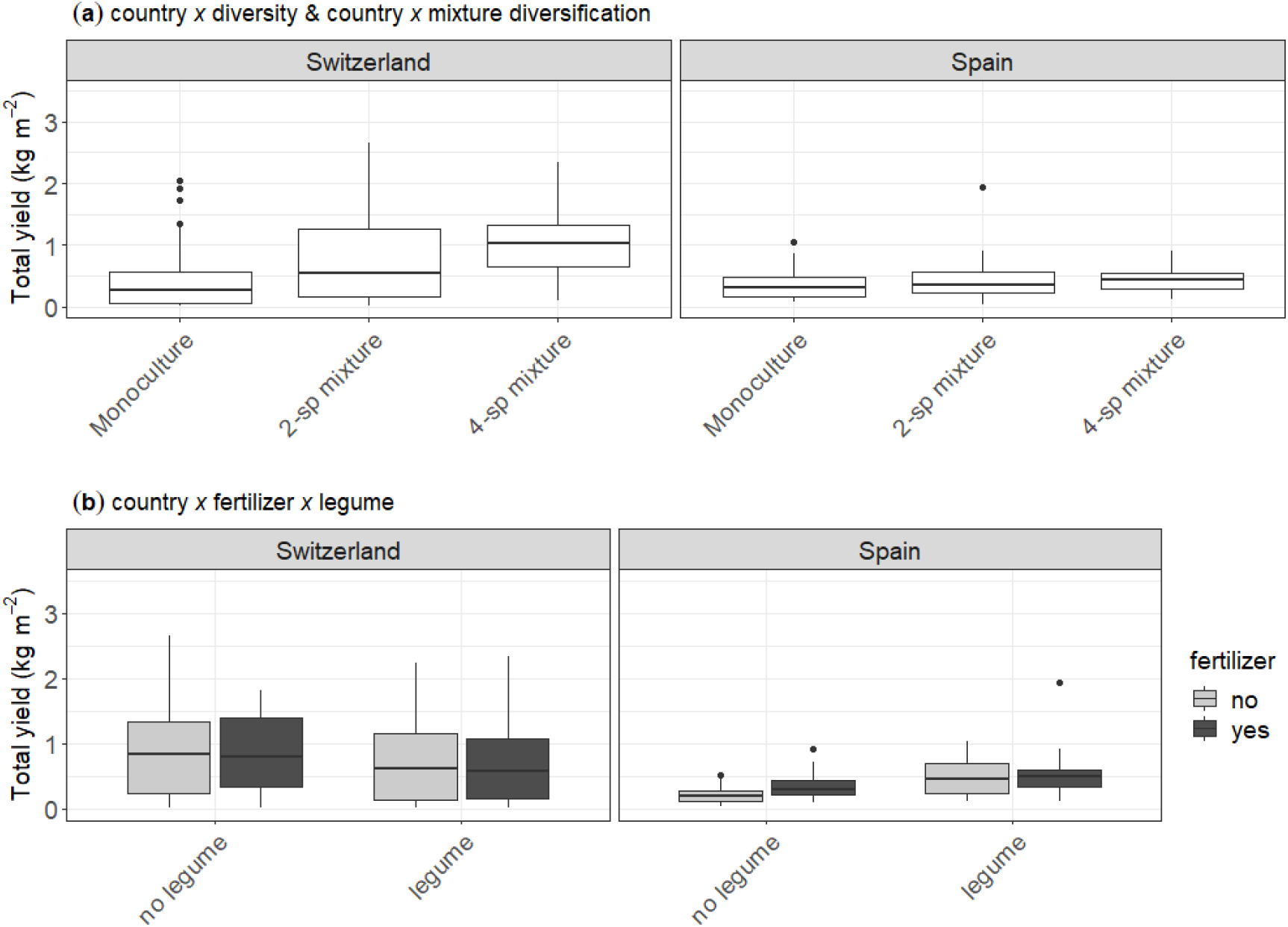
Community-level yields in kg dry weight per m^2^ visualizing the significant results from table 1. Differences in community-level yield between countries (**a**), diversity levels (**a**), mixture diversification (**a**), between the two-way interactions country × legume (**b**), country × diversity (**a**) and country × mixture diversification (**a**) and between the three-way interaction country × fertilizer × legume (**b**).

### 3.2. Environmental factors, plant traits and biodiversity effects

Neither environmental factors nor plant traits showed significant variation in response to the diversity treatments (Fig. 2, Table S4, S5). Reduced water input resulted in a significantly lower volumetric soil water content (VWC) in Spain. However, VWC did not vary significantly in response to fertilizer or diversity treatments (Fig. **2a**, Table S5). Both plant height and FPAR were significantly higher in Switzerland than in Spain, indicating that canopy closure was more complete and that vegetative growth was generally stronger in Switzerland than in Spain (Fig. **2b**,**e**) and in fertilized compared with unfertilized treatments (Table S5). SLA of crops was significantly higher in Switzerland compared with Spain and LDMC showed an opposite behavior, with higher values in Spain than in Switzerland (Fig. **2c**,**d**). Neither LDMC nor SLA responded significantly to fertilizer or diversity treatments. Leaf N and C:N ratio both indicate significantly higher leaf N levels in Switzerland than in Spain (Fig. **2f**,**g**, Table S4, S5). Complementarity effects were stronger in Switzerland than in Spain and stronger in 4-species than in 2-species mixtures in Switzerland (Fig. **2h**). Selection effects showed no response to any treatment factor (Fig. **2i**).

**Figure 2:**
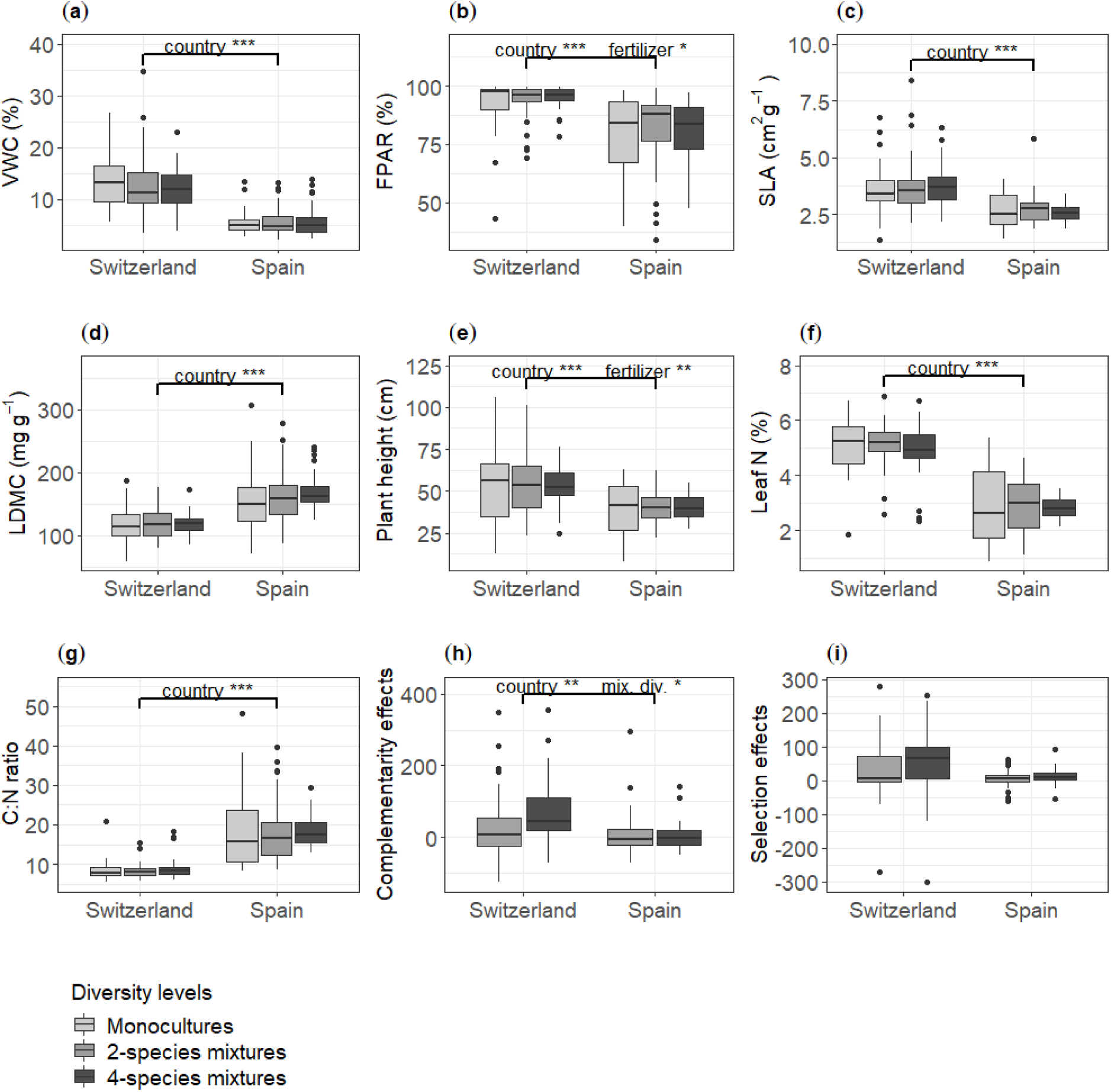
Community-level means for the environmental factors volumetric soil water content (VWC) (**a**) and absorption of photosynthetically active radiation (FPAR) (**b**), the plant traits specific leaf area (SLA) (**c**), leaf dry matter content (LDMC) (**d**), plant height (**e**), leaf N (**f**) and C:N ratio (**g**) and the biodiversity effects, divided into complementarity (**h**) and selection effects (**i**). Data are shown for both countries and separated by levels of diversity. Complementarity and selection effects are only available for mixtures. Brackets indicate significant differences between treatments and labels above brackets indicate which treatment (mix. div. = mixture diversification) was significant at α = 0.05 (* P < 0.05, ** P < 0.01, *** P < 0.001).

### 3.3 Structural equation model to explain community-level yields

The SEM showed a good fit to the data (d.f = 5, p = 0.572, CFI = 1) and explored the links between experimental treatment factors (country, fertilizer, mixture diversification), environmental factors (ΔFPAR, ΔVWC) and the effect of these on the relevant plant traits (ΔSLA; ΔLDMC; ΔC:N ratio, Δplant height) and finally linked these through selection and complementarity effects to yield differences between crop monocultures and mixtures (Δyield) (Fig. 3).

**Figure 3:**
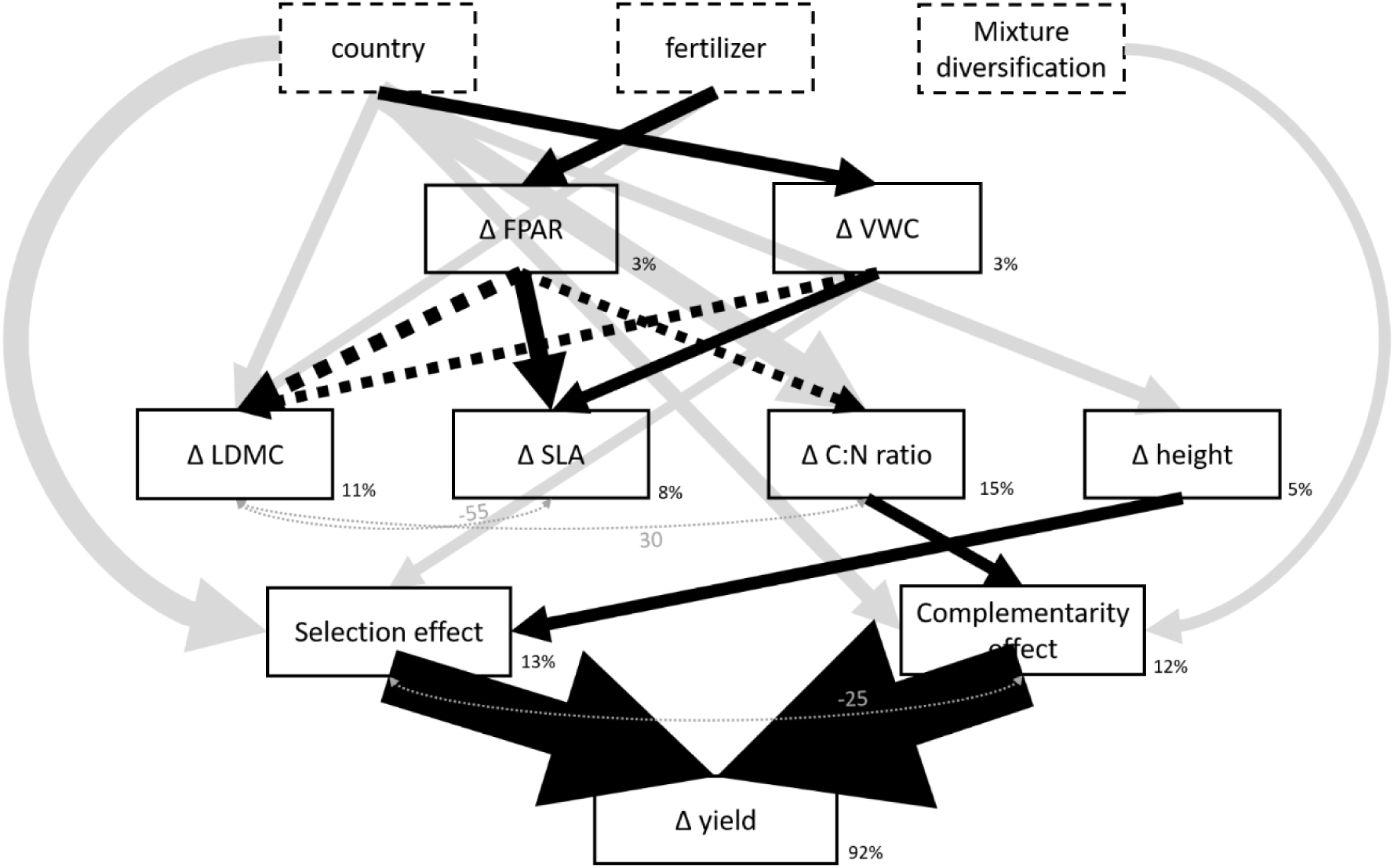
Mechanistic model showing the experimental treatment effects on the effects of environmental factors on plant traits and subsequently on biodiversity effects and differences in yield between crop mixtures and crop monocultures. Δ indicates differences between the respective measurements in mixtures compared with monocultures, thus positive Δ values indicate higher values in mixtures compared with monocultures and vice versa. Mean values for Δ values per country, fertilizer treatment and species number are given in table S6. Displayed black arrows show significant positive (solid) or negative (dashed) relationships (α = 0.05), grey arrows indicate direct effects of treatment factors on traits and yield. Arrow thickness indicates effect size based on standardized path coefficients. Numbers next to the variables indicate their explained variance (R^2^). Double-headed grey dashed arrows indicate significant correlations. Non-significant tested relationships are not shown.

The effect of complementarity on Δyield was 1.2 fold stronger than the selection effect (Fig. 3). Both biodiversity effects were negatively correlated with each other, thus when one effect increased, the other decreased. The selection effect was only related to Δheight, indicating that increasing plant height in mixtures compared to monocultures increased the selection effect (Fig. 3). We suspect that most of the selection effect is due to the tall-growing and high-yielding species *Chenopodium quinoa* dominating the communities and their yield, particularly in Switzerland (Fig. S3). The complementarity effect was positively related only to ΔC:N ratio. Increases in ΔC:N ratio indicate higher productivity per unit N in mixtures compared with monocultures, which increased the complementarity effect. Overall, the model explained 92% of yield differences between mixtures and monocultures (Fig. 3).

Plant traits, excluding plant height, were correlated among each other. ΔLDMC was strongly negatively correlated with ΔSLA and positively with ΔC:N ratio. This suggests that a higher water content in leaves of crop mixtures than crop monocultures (low ΔLDMC) is correlated to a higher leaf area per unit mass in crop mixtures than in crop monocultures (high ΔSLA) and to less leaf N in crop monocultures than in crop mixtures (low ΔC:N ratio) (Fig. 3).

ΔLDMC was negatively related to both environmental factors, ΔFPAR and ΔVWC, indicating that higher light interception by the crop canopy and higher soil water contents decreased LDMC, thus increasing leaf water contents in mixtures. The negative correlation between ΔC:N ratio and ΔFPAR implies that leaf N content in mixtures increased with increasing light interception. ΔSLA was positively related to both environmental factors, suggesting that increasing light interception and soil water content increased SLA, thus promoting higher leaf area per leaf mass. Based on standardized effect sizes, the effect of ΔFPAR on leaf traits was stronger than the effect of ΔVWC; specifically, the effect of ΔFPAR on ΔLDMC and ΔSLA was 1.4 and 1.5 fold stronger than the effect of ΔVWC, respectively. ΔVWC was positively correlated to the selection effect, with the effect of Δheight being 1.2 fold stronger than the effect of ΔVWC on the selection effect. Thus, higher soil water contents in mixtures compared with monocultures increased the selection effect in mixtures compared with monocultures (Fig. 3).

ΔFPAR varied in response to fertilizer treatment, with 10% higher values in unfertilized treatments, indicating that crop mixtures intercepted more light than crop monocultures and that this effect was stronger in unfertilized treatments (Table S6). ΔVWC varied significantly only among countries, with more negative values in Switzerland compared with Spain. This indicates that soils in crop monocultures had a higher water content than in crop mixtures and that this effect was more pronounced in Switzerland than in Spain. All plant traits, except ΔSLA varied significantly among countries; ΔLDMC was 163% higher and Δheight 207% higher in Spain than in Switzerland, while ΔC:N was 104% higher in Switzerland than in Spain (Fig. 3, Table S6). ΔLDMC also varied significantly among fertilization treatments, with 170% higher results in fertilized compared with unfertilized treatments. Both, selection and complementarity effects responded significantly to country. Complementarity effects were 540% higher in Switzerland than in Spain and selection effects were 350% higher in Switzerland than in Spain. Complementarity effect was the only variable responding to mixture diversification and complementarity effects were 110% higher in 4-than in 2-species mixtures. The effect of mixture diversification on complementarity effects was as strong as the effect from ΔC:N ratio on complementarity effects (Fig. 3, Table S6).

## 4. Discussion

Our study found increasing yields from crop monocultures to 2-to 4-species mixtures, in particular at the temperate site in Switzerland. Community-level yield did not respond to fertilizer treatments, but varied strongly between the two countries. Selection and complementarity effects were linked to plant height and C:N ratio, respectively, and explained 92% of the variation in community-level crop yield differences between mixtures and monocultures. While C:N ratio was linked to light use, plant height showed no link to environmental factors and only differed between countries. The effect of mixture diversification on crop yield was not mediated through any of the abiotic factors or plant traits measured in this study, but acted directly upon complementarity effects.

### Positive biodiversity–productivity relationships are context–dependent

In Switzerland community-level yield increased from monoculture to mixture and from 2-to 4-species mixtures, while the diversity effects on yield in Spain remained weak. The differences between the two countries were diverse and included differences in precipitation and irrigation, hours of sunshine, soil nutrients and soil texture. Light availability in Spain was presumably not limiting, but could have been an inhibiting factor. Lower SLA values in Spain indicate that the plants had less leaf area per dry leaf mass, which could be the plants’ effort to reduce leaf area exposed to high irradiance.

From the three growth-limiting resources we focused on in this study, soil water and N availability were the most promising to explain the missing positive diversity-productivity relationship in Spain. Soil water content in Spain was kept low by reducing irrigation to the amount needed for plant survival (see chapter 2.1) (mean VWC in Spain: 5.7 ± 0.4 %). Combined with a generally hotter climate in Spain, the crops were more prone to water stress. Crop yields in intercropping under drought conditions are expected to decrease (Coll et al. 2012). Also, positive diversity effects on crop water availability in intercropping remain contested and Brooker et al. (2015) suggest that these effects are limited to intercropping systems where at least one species has a low water demand. The crop species planted in this experiment were not adapted to the dry conditions in Spain, since they were Swiss cultivars bred for use under temperate climatic conditions. A further explanation for the absence of a positive diversity-productivity relationship in Spain can be the increased allocation of C to belowground productivity in response to dry conditions. In our study we were interested in positive productivity effects on crop yield, hence the focus on above-ground biomass. However, increased belowground investment can lead to a decrease in aboveground productivity while maintaining overall community productivity (Kahmen et al. 2005).

Besides arid conditions in Spain, low soil fertility could also have reduced biodiversity effects on productivity, as observed before in an agricultural experiment (Fridley 2003). The increase in community-level yields in response to legume presence in Spain indicates that N sparing by the legumes could increase leaf N content for neighboring crops. Since the N sparing effect was only observed in unfertilized communities in Spain, we suggest that fertilizer addition reduced the formation of nodules during early legume growth, inhibiting N-fixation during later growth stages (Naudin et al. 2011). Also, available soil water content is an important parameter controlling N_2_-fixation of legumes, either directly by influencing nodulation or indirectly by reducing plant growth and thus N_2_-fixation (Sprent and Minchin 1983). Therefore, we suggest that drier conditions in Spain might have limited plant growth directly and also inhibited N_2_-fixation, further reducing N availability for crops.

### Complementarity effects increased yields in mixtures compared with monocultures

In our study, complementarity effects were shown to drive positive biodiversity–productivity relationships in Switzerland. The effect of complementarity was mainly linked to changes in C:N ratio between mixtures and monocultures, implying that crops grown in mixtures were producing more biomass per unit leaf N than crops in monoculture. This increase in efficiency could partially explain the higher productivity often observed at higher diversity (Fargione et al. 2007). The C:N ratio was dependent on the fraction of intercepted light but not on soil water contents. The negative link between C:N ratio and FPAR indicated that higher light interception was linked to higher leaf N content per unit biomass. Thus, crops responded to a more complete canopy cover and thus lower light access by increasing their SLA (Fig. 3) and leaf N content to have a larger photosynthetically active leaf area. High leaf N and high SLA are a common plant response to lower light conditions (Lambers et al. 1998; Reich et al. 1998; Evans and Poorter 2001; Funk et al. 2017). However in Switzerland, at higher mixture diversification (i.e. in 4-species mixtures) the relation between FPAR and C:N ratio became positive (table S6), indicating that with increasing diversification a shift in N use efficiency occurred. Thus, 4-species mixtures in Switzerland produced more yield per unit N than 2-species mixtures or monocultures. Increasing N efficiency in more diverse communities has been observed before in semi-natural (van Ruijven and Berendse 2005; Fargione et al. 2007) but, to the best of our knowledge, not in intercropping systems.

Complementarity effects were the only variable responding to mixture diversification, with an increase of complementarity effects in 4-over 2-species mixtures. Strikingly, mixture diversification did not affect any of the environmental factors or plant traits measured in this study. This suggests that the plant traits measured in this study were not able to describe how mixture diversification affects complementarity. We suspect belowground processes driving yield increases from 2- to 4-species mixtures or that dynamics of nutrients other than N, or reduced impacts of pests, were possibly playing a role. Concerning possible belowground processes, other studies observed that the presence of a legume can increase N or P availability in the soil surrounding its roots (Temperton et al. 2007; Zhang et al. 2019). In our study, this observation could potentially be important, as all 4-species mixtures contained one leguminous crop, while not all 2-species mixtures did so. As an alternative to not measuring the appropriate traits, it could also be that not enough traits were measured to fully capture niche differences. As suggested by earlier studies, when linking plant traits to biodiversity effects, a large range of traits is required to explain niche differences (Kraft et al. 2015; Cadotte 2017).

Although we did see facilitative processes in Spain, especially the N sparing effect of the legumes in unfertilized plots, we propose that facilitative processes were not strong enough to compensate for the difficult growing conditions in Spain.

### Selection effects due to Chenopodium quinoa

In this study, selection effects had a similarly strong effect on yield differences between mixtures and monocultures as had complementarity effects. While it is often assumed that positive biodiversity– productivity relationships are driven mainly by niche differentiation (Cardinale 2013), we show here that selection effects are nearly as important. The selection effect was linked to plant height, thus selection effects increased with increasing plant height in mixtures compared to monocultures. A relation between plant height and selection effects has been observed before (Cadotte 2017) and is probably due to plant height being related to competition for light, where taller plants outcompete shorter plants (Westoby 1998).

Selection effects were significantly higher in Switzerland than in Spain. Combined with the link to plant height, we concluded that one highly productive and tall-growing species was causing this effect *Chenopodium quinoa* (Fig. S3). It has been observed before that *C. quinoa* was highly competitive (Buckland 2016) and that abundant irrigation could increase its yields (Walters et al. 2016). Thus, the selection effect was considerably less important in Spain than in Switzerland.

## 5. Conclusion

Our study showed that crop productivity increased with diversity under temperate conditions but only weakly when crops were grown under semiarid conditions with limited availability of water and strong irradiance. Increases in productivity in mixtures compared to monocultures were caused to almost the same extent by complementarity and selection effects. Selection and complementarity effects were explained by different plant trait syndromes. The selection effect was maximized in plots with tall plants and was probably caused by one single species, *Chenopodium quinoa*, which was highly productive and tall-growing. Complementarity effects were linked to an increased C capture per unit N, thus indicating that crops in mixtures were more efficient. However, complementarity effects were also stronger in 4-compared with 2-species mixtures and this link was not mediated through any of the measured plant traits, suggesting that other ecological processes must have been responsible for the positive diversity effect on yield beyond two-species mixtures. This finding suggests that the drivers of diversity effects from monocultures to mixtures are not the same as from 2- to 4-species mixtures and should therefore be targeted specifically in future studies. Nevertheless, by identifying links between relevant plant traits and positive biodiversity–productivity relationships through selection and complementarity effects, this study helps pave the way to select crop species for mixtures that optimize sampling and complementarity effects.

## Acknowledgments

We are grateful to Elisa Pizarro Carbonell, Carlos Barriga Cabanillas and Anja Schmutz for their help with the field experiment, and Johan Six for comments on the experimental design. We also thank the Aprisco de Las Corchuelas and the University of Zurich for allowing us to use their experimental gardens. The study was funded by the Swiss National Science Foundation (PP00P3_170645).

## Author contribution

N.E. and C.S. conceived the study with input from L.S. and R.W.B. N.E., L.S. and C.S. collected the data; N.E. assembled and analyzed the data with the help of C.S.; N.E. and C.S. wrote the paper. All authors discussed data analyses and results and revised the manuscript.

## Notes

### Competing Interest Statement

The authors have declared no competing interest.

