## supplementinfo for "Using plant traits to understand the contribution of biodiversity effects to community productivity in an agricultural system"

### *New Phytologist* Supporting Information

Article acceptance date: Click here to enter a date.

The following Supporting Information is available for this article:

**Fig. S1** *Cross-reference for N and C content analysis on LECO and MS.*

Cross-reference of 8 samples for N and C leaf content analysis on LECO (LECO CHN628C elemental analyzer [Leco Co., St. Joseph, USA]) and MS (PDZ Europa 20-20 isotope ratio mass spectrometer linked to a PDZ Europa ANCA-GSL elemental analyzer [Sercon Ltd., Cheshire, UK]). Samples were analyzed on either device, depending on the available leaf dry mass. Samples with less than 100 mg (505 samples) available leaf dry mass were measured on MS and samples with more than 100 mg (400 samples) available leaf dry mass were measured on LECO.

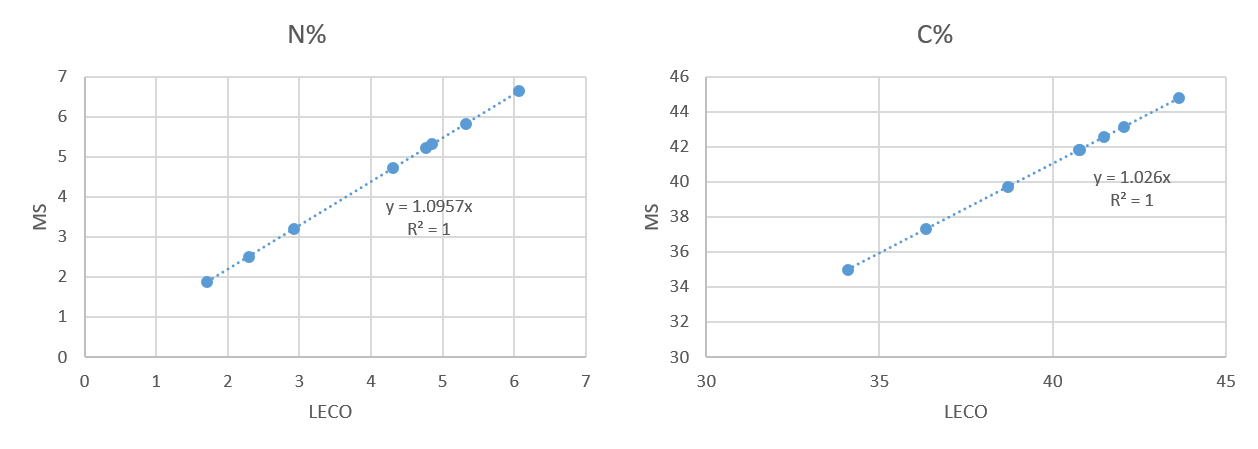

**Fig. S2** *A priori structural model for the relationship of variables analyzed with the structural equation model.*

Our a priori model related environmental factors to plant traits to selection and complementarity effects to yield differences between mixtures and monocultures. Also, the dependence on any of these to the treatment factors were analysed.

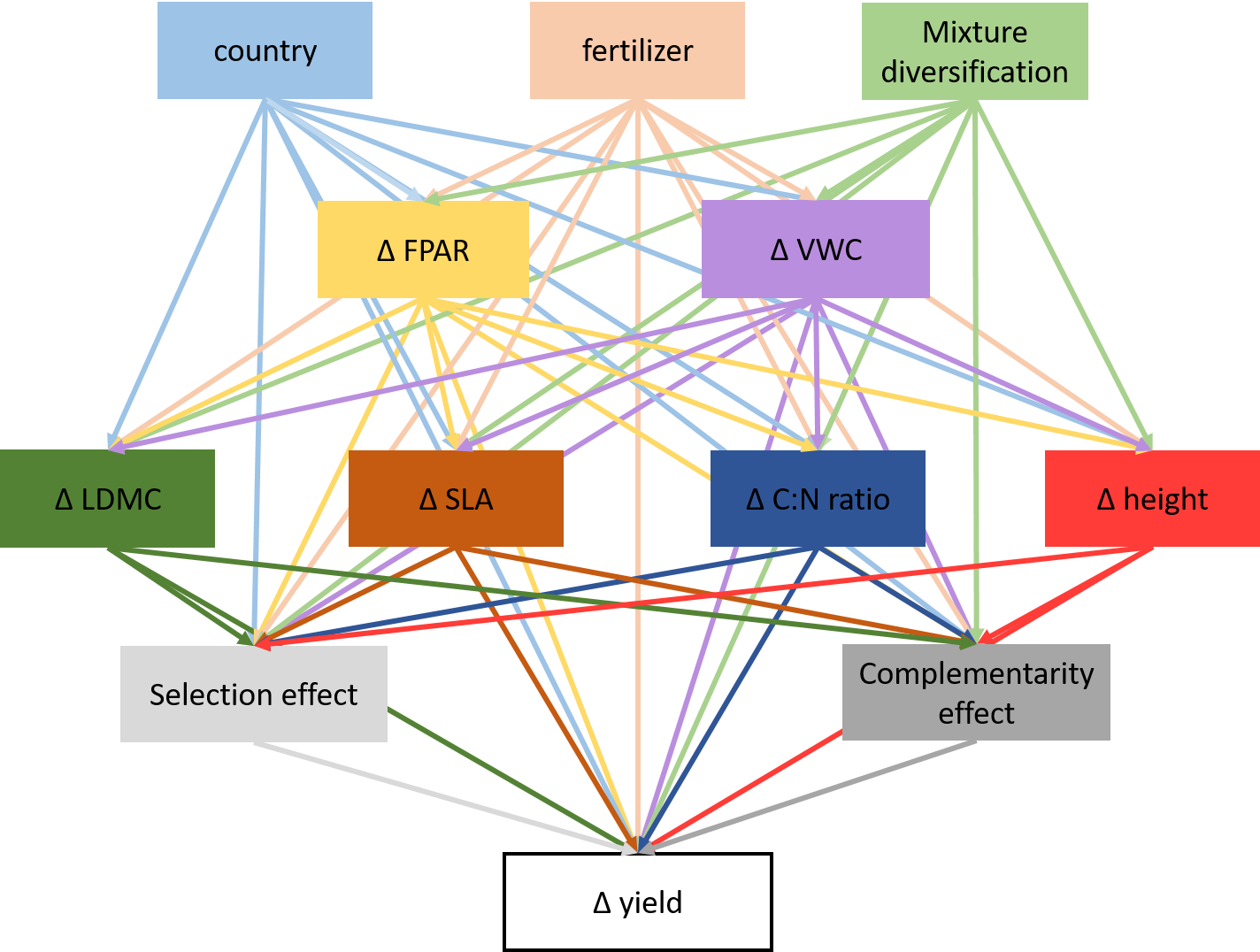

**Fig. S3** *ΔYield (A) and Δplant height (B) of the eight different species in both countries.*

ΔYield (A) and Δplant height (B) of the eight different species (Ave = Avena sativa, Cam = Camelina sativa, Cor = Coriandrum sativum, Len = Lens culinaris, Lin = Linum usitatissimum, Lup = Lupinus angustifolius, Qui = Chenopodium quinoa, Tri = Triticum aestivum) separated by country. Δ are calculated as value in mixture – mean value in monoculture. To make mixtures comparable with monocultures, we multiplied the value in mixture with the number of species. The red dotted line indicates 0. ΔValues above 0 indicate higher values in mixtures compared with monocultures and Δvalues below 0 indicate lower values in mixtures compared with monocultures.

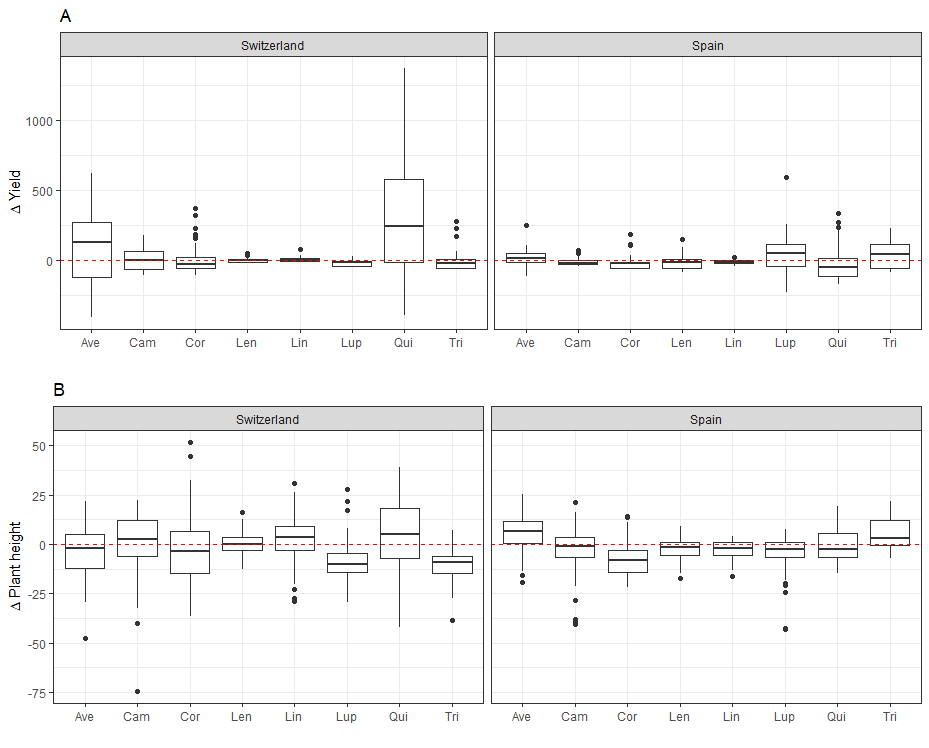

**Table S1** *List of crop species phylogenetic groups, cultivar, seed supplier and their sowing densities****.***

| Species | Phylogenetic group | Cultivar | Supplier | Sowing density (seeds / m^2^) |
| --- | --- | --- | --- | --- |
| *Triticum aestivum* | Cereal | Fiorina | DSP, Delley | 400 |
| *Avena sativa* | Cereal | Canyon | Sativa Rheinau | 400 |
| *Lens culinaris* | Legume | Anicia | Agroscope, Reckenholz | 160 |
| *Lupinus angustifolius* | Legume | Boregine | Aspenhof, Wilchingen | 160 |
| *Camelina sativa* | Superrosid | n.a. | Zollinger Samen, Les Evouettes | 592 |
| *Linum usitatissimum* | Superrosid | Lirina | Sativa Rheinau | 592 |
| *Chenopodium quinoa* | Superasterid | n.a. | Artha Samen, Münsingen | 240 |
| *Coriandrum sativum* | Superasterid | Indian | Zollinger Samen, Les Evouettes | 240 |

**Table S2** *Crop growth duration in mean days from sowing (± SD) to harvest for both countries and all eight species.*

| Species | Days after sowing | |
| --- | --- | --- |
|  | Switzerland | Spain |
| *Avena sativa* | 120 ± 6.2 | 146 ± 6.2 |
| *Triticum aestivum* | 106 ± 9.9 | 146 ± 1.3 |
| *Lens culinaris* | 128 ± 16.2 | 146 ± 1 |
| *Lupinus angustifolius* | 126 ± 15.7 | 134 ± 1.7 |
| *Camelina sativa* | 98 ± 1 | 153 ± 4.4 |
| *Linum usitatissimum* | 136 ± 3.1 | 155 ± 3.1 |
| *Coriandrum sativum* | 123 ± 10.7 | 152 ± 5.5 |
| *Chenopodium quinoa* | 142 ± 7.8 | 187 ± 6.4 |

**Table S3** *Tukey post-hoc test of linear mixed effects model in table 1.*

Tukey post-hoc test of linear mixed effects model testing effects of the different treatments on community-level yield (results in table 1). P-values in bold are significant at α = 0.05 (* P < 0.05, ** P < 0.01, *** P < 0.001). CH = Switzerland, ES = Spain, mix = mixture, mono = monoculture, 2 = 2-species mixture, 4 = 4-species mixture. Only significant interactions are shown for the threeway-interaction country x fertilized x legume.

| contrast | estimate | SE | df | t.ratio | p.value |
| --- | --- | --- | --- | --- | --- |
| **~ country x diversity** | | | | | |
| **CH,mix-ES,mix** | 1.02 | 0.11 | 33.7 | 9.21 | **<.0001***** |
| **CH,mix-CH,mono** | 1.59 | 0.30 | 54 | 5.25 | **<.0001***** |
| CH,mix-ES,mono | 0.82 | 0.3 | 52.5 | 2.72 | 0.042 |
| ES,mix-CH,mono | 0.57 | 0.3 | 52.8 | 1.9 | 0.243 |
| ES,mix-ES,mono | -0.2 | 0.31 | 55.9 | -0.65 | 0.915 |
| **CH,mono-ES,mono** | -0.77 | 0.17 | 103.4 | -4.49 | **0.0001**** |
| **~ country x mixture diversification** | | | | | |
| **2,mix,CH-4,mix,CH** | -1.36 | 0.27 | 53.5 | -5.12 | **0.0001**** |
| **2,mix,CH-mono,CH** | 0.91 | 0.29 | 53.1 | 3.09 | **0.036*** |
| 2,mix,CH-2,mix,ES | 0.25 | 0.12 | 31.4 | 2.2 | 0.268 |
| 2,mix,CH-4,mix,ES | 0.42 | 0.27 | 58.6 | 1.56 | 0.63 |
| 2,mix,CH-mono,ES | 0.14 | 0.3 | 54 | 0.47 | 0.997 |
| **4,mix,CH-mono,CH** | 2.27 | 0.36 | 55.5 | 6.25 | **<.0001***** |
| **4,mix,CH-2,mix,ES** | 1.61 | 0.27 | 59.2 | 5.96 | **<.0001***** |
| **4,mix,CH-4,mix,ES** | 1.78 | 0.17 | 99 | 10.45 | **<.0001***** |
| **4,mix,CH-mono,ES** | 1.5 | 0.36 | 52.9 | 4.18 | **0.002**** |
| mono,CH-2,mix,ES | -0.65 | 0.3 | 54.1 | -2.21 | 0.25 |
| mono,CH-4,mix,ES | -0.49 | 0.36 | 52.6 | -1.36 | 0.751 |
| **mono,CH-mono,ES** | -0.77 | 0.17 | 103.4 | -4.49 | **0.0003***** |
| 2,mix,ES-4,mix,ES | 0.17 | 0.26 | 49.9 | 0.63 | 0.988 |
| 2,mix,ES-mono,ES | -0.12 | 0.3 | 55.3 | -0.39 | 0.999 |
| 4,mix,ES-mono,ES | -0.28 | 0.36 | 56.2 | -0.77 | 0.971 |
| **~ country x legume** | | | | | |
| **CH,no - ES,no** | 1.158 | 0.132 | 154.3 | 8.761 | **<.0001***** |
| **CH,no - CH,yes** | 1.274 | 0.256 | 54.2 | 4.985 | **<.0001***** |
| CH,no - ES,yes | 0.364 | 0.259 | 56.2 | 1.406 | 0.501 |
| ES,no - CH,yes | 0.116 | 0.261 | 57.9 | 0.445 | 0.9704 |
| **ES,no - ES,yes** | -0.794 | 0.259 | 56.4 | -3.068 | 0.0169 |
| **CH,yes - ES,yes** | -0.91 | 0.131 | 145.3 | -6.954 | **<.0001***** |
| **~ country x fertilizer x legume** | | | | | |
| **CH,no,no - ES,no,no** | 1.4574 | 0.186 | 150 | 7.851 | **<.0001** |
| **CH,no,no - ES,yes,no** | 0.8878 | 0.186 | 155.7 | 4.768 | **0.0001** |
| **CH,no,no - CH,no,yes** | 1.4651 | 0.284 | 80.1 | 5.16 | **<.0001** |
| **ES,no,no - CH,yes,no** | -1.429 | 0.186 | 149.1 | -7.676 | **<.0001** |
| **ES,no,no - ES,no,yes** | -1.1006 | 0.291 | 85.5 | -3.785 | **0.0066** |
| **CH,yes,no - CH,no,yes** | 1.4367 | 0.289 | 83 | 4.973 | **0.0001** |
| **CH,yes,no - CH,yes,yes** | 1.0837 | 0.283 | 79.2 | 3.831 | **0.0059** |
| **CH,no,yes - ES,no,yes** | -1.1083 | 0.185 | 150.6 | -5.979 | **<.0001** |
| **CH,no,yes - ES,yes,yes** | -1.0656 | 0.184 | 142.6 | -5.796 | **<.0001** |
| **ES,no,yes - CH,yes,yes** | 0.7552 | 0.185 | 145.3 | 4.073 | **0.0019** |
| **CH,yes,yes - ES,yes,yes** | -0.7125 | 0.185 | 137.9 | -3.862 | **0.0042** |

**Table S4** *Means ± standard error for environmental factors and plant traits in both countries.*

Mean ± SE for environmental factors and plant traits in Switzerland (CH) and Spain (ES), with and without application of fertilizer in monocultures (mono), 2-species mixtures (2-sp-mix) and 4-species mixtures (4-sp-mix)). Measurements were taken at the time of flowering.

| country | fertilized | diversity | VWC  [%] | FPAR  [%] | SLA  [cm^2^ g^-1^] | LDMC  [mg g^-1^] | plant height [cm] | leaf N  [%] | C:N ratio |
| --- | --- | --- | --- | --- | --- | --- | --- | --- | --- |
| CH | no | mono | 13.3 ± 1.56 | 90.8 ± 3.46 | 3.69 ± 0.35 | 116.9 ± 7.63 | 54 ± 6.24 | 5.3 ± 0.22 | 7.8 ± 0.39 |
|  |  | 2-sp-mix | 11.9 ± 0.71 | 93.4 ± 1.07 | 3.65 ± 0.15 | 121.8 ± 3.56 | 55.3 ± 3.04 | 5.24 ± 0.09 | 7.96 ± 0.18 |
|  |  | 4-sp-mix | 11.5 ± 0.7 | 95.2 ± 0.99 | 3.98 ± 0.19 | 116.2 ± 3.37 | 52.9 ± 2.29 | 5.17 ± 0.13 | 8.05 ± 0.23 |
|  | yes | mono | 14.3 ± 1.16 | 93.2 ± 2.35 | 3.44 ± 0.23 | 119.2 ± 7.76 | 55.5 ± 5.6 | 4.93 ± 0.28 | 8.8 ± 0.9 |
|  |  | 2-sp-mix | 13.6 ± 0.92 | 95.6 ± 0.79 | 3.7 ± 0.17 | 116.4 ± 3.49 | 53.6 ± 2.44 | 5.1 ± 0.11 | 8.19 ± 0.28 |
|  |  | 4-sp-mix | 12.7 ± 0.96 | 95.7 ± 0.74 | 3.48 ± 0.12 | 120.6 ± 2.68 | 55.1 ± 2.35 | 4.73 ± 0.25 | 9.56 ± 0.77 |
| ES | no | mono | 5.45 ± 0.74 | 74.7 ± 4.86 | 2.49 ± 0.17 | 166 ± 14.6 | 37.4 ± 3.84 | 3.08 ± 0.34 | 17.7 ± 2.64 |
|  |  | 2-sp-mix | 5.87 ± 0.49 | 81.1 ± 2.55 | 2.63 ± 0.07 | 163.9 ± 5.84 | 37.1 ± 1.57 | 2.9 ± 0.14 | 17.3 ± 1.11 |
|  |  | 4-sp-mix | 5.91 ± 0.69 | 80.2 ± 3.1 | 2.59 ± 0.07 | 173.7 ± 5.46 | 35.2 ± 0.99 | 2.93 ± 0.08 | 17.6 ± 0.81 |
|  | yes | mono | 6.15 ± 0.78 | 83.4 ± 3.37 | 2.78 ± 0.2 | 143.5 ± 8.36 | 44 ± 3.74 | 2.68 ± 0.36 | 19 ± 2.19 |
|  |  | 2-sp-mix | 5.28 ± 0.3 | 84.5 ± 1.81 | 2.87 ± 0.12 | 157.6 ± 6.37 | 44.3 ± 1.61 | 2.92 ± 0.16 | 17.3 ± 1.08 |
|  |  | 4-sp-mix | 5.85 ± 0.56 | 80.3 ± 2.56 | 2.57 ± 0.07 | 163.1 ± 5.69 | 44.4 ± 1.2 | 2.69 ± 0.07 | 18.6 ± 0.67 |

**Table S5** *Results of mixed effects ANOVA testing effects of treatments, legume presence, diversity and mixture diversification on community-level yield.*

Results of mixed effects ANOVA testing effects of the different treatments (country, fertilizer, legume, diversity (monocultures vs. mixtures) and mixture diversification (2- vs. 4-species mixtures) on community-level yield. on environmental variables (VWC, FPAR), plant traits (SLA, LDMC, plant height, leaf N, C:N ratio) and biodiversity effects (complementarity & selection effects). Df: degrees of freedom, SS: Sum of squares, MS: mean of squares, F-value: variance ratio, P: error probability. P-values in bold are significant at α = 0.05 (* P < 0.05, ** P < 0.01, *** P < 0.001). n = 315

| **VWC** | | | | | | | |
| --- | --- | --- | --- | --- | --- | --- | --- |
|  | SS | MS | numDF | denDF | F-value | P |  |
| country | 24.575 | 24.575 | 1 | 36.79 | 173.39 | **1.71E-15** | *** |
| fertilizer | 0.217 | 0.217 | 1 | 36.66 | 1.53 | 0.22 |  |
| diversity | 0.078 | 0.078 | 1 | 40.99 | 0.55 | 0.46 |  |
| mix. diversification | 0.002 | 0.002 | 1 | 48.29 | 0.02 | 0.90 |  |
| country × fertilizer | 0.051 | 0.051 | 1 | 36.75 | 0.36 | 0.55 |  |
| country × diversity | 0.169 | 0.169 | 1 | 260.51 | 1.19 | 0.28 |  |
| country × mix. diversification | 0.036 | 0.036 | 1 | 271.55 | 0.25 | 0.62 |  |
| fertilizer × diversity | 0.022 | 0.022 | 1 | 256.67 | 0.15 | 0.70 |  |
| fertilizer × mix. diversification | 0.037 | 0.037 | 1 | 262.47 | 0.26 | 0.61 |  |
| country × fertilizer × diversity | 0.007 | 0.007 | 1 | 260.21 | 0.05 | 0.83 |  |
| country × fertilizer × mix. diversification | 0.047 | 0.047 | 1 | 267.57 | 0.33 | 0.57 |  |
| **FPAR** | | | | | | | |
| country | 1.388 | 1.388 | 1 | 36.20 | 61.57 | **2.53E-09** | *** |
| fertilizer | 0.120 | 0.120 | 1 | 36.05 | 5.31 | **0.03** | * |
| diversity | 0.025 | 0.025 | 1 | 40.69 | 1.09 | 0.30 |  |
| mix. diversification | 0.004 | 0.004 | 1 | 47.81 | 0.19 | 0.67 |  |
| country × fertilizer | 0.033 | 0.033 | 1 | 36.12 | 1.46 | 0.23 |  |
| country × diversity | 0.005 | 0.005 | 1 | 263.46 | 0.22 | 0.64 |  |
| country × mix. diversification | 0.027 | 0.027 | 1 | 274.26 | 1.22 | 0.27 |  |
| fertilizer × diversity | 0.064 | 0.064 | 1 | 259.89 | 2.83 | 0.09 |  |
| fertilizer × mix. diversification | 0.017 | 0.017 | 1 | 265.31 | 0.74 | 0.39 |  |
| country × fertilizer × diversity | 0.020 | 0.020 | 1 | 263.23 | 0.87 | 0.35 |  |
| country × fertilizer × mix. diversification | 0.001 | 0.001 | 1 | 269.73 | 0.06 | 0.81 |  |
| **SLA** | | | | | | | |
| country | 4.327 | 4.327 | 1 | 26.43 | 148.60 | **2.35E-12** | *** |
| fertilizer | 0.008 | 0.008 | 1 | 26.20 | 0.26 | 0.61 |  |
| diversity | 0.009 | 0.009 | 1 | 43.80 | 0.32 | 0.58 |  |
| mix. diversification | 0.000 | 0.000 | 1 | 46.00 | 0.00 | 0.95 |  |
| country × fertilizer | 0.139 | 0.139 | 1 | 26.38 | 4.77 | **0.04** | * |
| country × diversity | 0.072 | 0.072 | 1 | 251.59 | 2.47 | 0.12 |  |
| country × mix. diversification | 0.072 | 0.072 | 1 | 255.23 | 2.48 | 0.12 |  |
| fertilizer × diversity | 0.044 | 0.044 | 1 | 249.57 | 1.51 | 0.22 |  |
| fertilizer × mix. diversification | 0.059 | 0.059 | 1 | 251.88 | 2.04 | 0.15 |  |
| country × fertilizer × diversity | 0.001 | 0.001 | 1 | 252.99 | 0.05 | 0.82 |  |
| country × fertilizer × mix. diversification | 0.031 | 0.031 | 1 | 255.58 | 1.07 | 0.30 |  |
| **LDMC** | | | | | | | |
| country | 3.364 | 3.364 | 1 | 21.42 | 155.38 | **2.76E-11** | *** |
| fertilizer | 0.021 | 0.021 | 1 | 21.24 | 0.98 | 0.33 |  |
| diversity | 0.019 | 0.019 | 1 | 44.16 | 0.88 | 0.35 |  |
| mix. diversification | 0.004 | 0.004 | 1 | 46.26 | 0.19 | 0.66 |  |
| country × fertilizer | 0.057 | 0.057 | 1 | 21.39 | 2.63 | 0.12 |  |
| country × diversity | 0.075 | 0.075 | 1 | 246.90 | 3.47 | 0.06 |  |
| country × mix. diversification | 0.017 | 0.017 | 1 | 250.12 | 0.76 | 0.38 |  |
| fertilizer × diversity | 0.005 | 0.005 | 1 | 244.82 | 0.22 | 0.64 |  |
| fertilizer × mix. diversification | 0.001 | 0.001 | 1 | 247.31 | 0.06 | 0.80 |  |
| country × fertilizer × diversity | 0.001 | 0.001 | 1 | 248.21 | 0.02 | 0.88 |  |
| country × fertilizer × mix. diversification | 0.036 | 0.036 | 1 | 250.73 | 1.67 | 0.20 |  |
| **Plant height** | | | | | | | |
| country | 3.682 | 3.682 | 1 | 24.85 | 62.67 | **2.97E-08** | *** |
| fertilizer | 0.687 | 0.687 | 1 | 24.66 | 11.69 | **0.00** | ** |
| diversity | 0.020 | 0.020 | 1 | 44.86 | 0.34 | 0.56 |  |
| mix. diversification | 0.001 | 0.001 | 1 | 47.81 | 0.02 | 0.88 |  |
| country × fertilizer | 0.356 | 0.356 | 1 | 24.80 | 6.05 | **0.02** | * |
| country × diversity | 0.017 | 0.017 | 1 | 254.86 | 0.28 | 0.60 |  |
| country × mix. diversification | 0.045 | 0.045 | 1 | 259.50 | 0.76 | 0.39 |  |
| fertilizer × diversity | 0.004 | 0.004 | 1 | 252.59 | 0.08 | 0.78 |  |
| fertilizer × mix. diversification | 0.042 | 0.042 | 1 | 255.15 | 0.71 | 0.40 |  |
| country × fertilizer × diversity | 0.000 | 0.000 | 1 | 256.32 | 0.00 | 0.97 |  |
| country × fertilizer × mix. diversification | 0.018 | 0.018 | 1 | 259.40 | 0.31 | 0.58 |  |
| **Leaf N** | | | | | | | |
| country | 5.732 | 5.732 | 1 | 20.41 | 123.18 | **4.21E-10** | *** |
| fertilizer | 0.122 | 0.122 | 1 | 20.32 | 2.63 | 0.12 |  |
| diversity | 0.011 | 0.011 | 1 | 45.02 | 0.24 | 0.62 |  |
| mix. diversification | 0.008 | 0.008 | 1 | 49.03 | 0.16 | 0.69 |  |
| country × fertilizer | 0.003 | 0.003 | 1 | 20.38 | 0.06 | 0.81 |  |
| country × diversity | 0.098 | 0.098 | 1 | 244.08 | 2.12 | 0.15 |  |
| country × mix. diversification | 0.071 | 0.071 | 1 | 249.82 | 1.53 | 0.22 |  |
| fertilizer × diversity | 0.010 | 0.010 | 1 | 242.09 | 0.21 | 0.65 |  |
| fertilizer × mix. diversification | 0.025 | 0.025 | 1 | 245.94 | 0.54 | 0.46 |  |
| country × fertilizer × diversity | 0.007 | 0.007 | 1 | 245.37 | 0.15 | 0.70 |  |
| country × fertilizer × mix. diversification | 0.000 | 0.000 | 1 | 249.67 | 0.00 | 0.95 |  |
| **C:N ratio** | | | | | | | |
| country | 7.808 | 7.808 | 1 | 20.27 | 164.05 | **3.54E-11** | *** |
| fertilizer | 0.089 | 0.089 | 1 | 20.17 | 1.87 | 0.19 |  |
| diversity | 0.037 | 0.037 | 1 | 44.61 | 0.78 | 0.38 |  |
| mix. diversification | 0.093 | 0.093 | 1 | 48.45 | 1.96 | 0.17 |  |
| country × fertilizer | 0.015 | 0.015 | 1 | 20.23 | 0.31 | 0.59 |  |
| country × diversity | 0.008 | 0.008 | 1 | 243.58 | 0.17 | 0.68 |  |
| country × mix. diversification | 0.003 | 0.003 | 1 | 249.15 | 0.07 | 0.79 |  |
| fertilizer × diversity | 0.003 | 0.003 | 1 | 241.64 | 0.06 | 0.81 |  |
| fertilizer × mix. diversification | 0.037 | 0.037 | 1 | 245.41 | 0.78 | 0.38 |  |
| country × fertilizer × diversity | 0.014 | 0.014 | 1 | 244.90 | 0.29 | 0.59 |  |
| country × fertilizer × mix. diversification | 0.009 | 0.009 | 1 | 249.12 | 0.19 | 0.66 |  |
| **Complementarity effects** | | | | | | | |
| country | 44970 | 44970 | 1 | 30.01 | 11.10 | **0.00** | ** |
| fertilizer | 1727 | 1727 | 1 | 29.43 | 0.43 | 0.52 |  |
| mix. diversification | 17446 | 17446 | 1 | 29.06 | 4.31 | **0.05** | * |
| country × fertilizer | 429 | 429 | 1 | 29.49 | 0.11 | 0.75 |  |
| country × mix. diversification | 15680 | 15680 | 1 | 193.06 | 3.87 | 0.05 |  |
| fertilizer × mix. diversification | 2238 | 2238 | 1 | 165.00 | 0.55 | 0.46 |  |
| country × fertilizer × mix. diversification | 517 | 517 | 1 | 168.99 | 0.13 | 0.72 |  |
| **Selection effects** | | | | | | | |
| country | 4314.5 | 4314.5 | 1 | 188.89 | 1.57 | 0.21 |  |
| fertilizer | 5242.1 | 5242.1 | 1 | 163.06 | 1.91 | 0.17 |  |
| mix. diversification | 2575.6 | 2575.6 | 1 | 34.77 | 0.94 | 0.34 |  |
| country × fertilizer | 116 | 116 | 1 | 165.83 | 0.04 | 0.84 |  |
| country × mix. diversification | 278.5 | 278.5 | 1 | 188.89 | 0.10 | 0.75 |  |
| fertilizer × mix. diversification | 87.7 | 87.7 | 1 | 163.06 | 0.03 | 0.86 |  |
| country × fertilizer × mix. diversification | 357.9 | 357.9 | 1 | 165.83 | 0.13 | 0.72 |  |

**Table S6** *Mean values ± standard errors for Δenvironmental factors, Δplant traits, Δyield, complementarity (CE) and selection (SE) effects.*

Differences of plant traits, yield and environmental factors between community-level means in mixtures compared to community-level means in monocultures. Δ values were calculated according to formula 1-3 (see methods). Negative Δ values indicate higher community-level means in monocultures compared with mixtures.

| country | CH | | | | ES | | | |
| --- | --- | --- | --- | --- | --- | --- | --- | --- |
| fertilized | no | | yes | | no | | yes | |
| species nr. | 2 | 4 | 2 | 4 | 2 | 4 | 2 | 4 |
| ΔFPAR | 2.5 ± 1.34 | 4.44 ± 1.11 | 2.39 ± 0.67 | 3.18 ± 0.75 | 6.52 ± 2.2 | 5.35 ± 3.06 | 1.03 ± 1.95 | -3.24 ± 2.41 |
| ΔVWC | -1.4 ± 0.82 | -1.75 ± 0.68 | -0.71 ± 0.77 | -1.65 ± 0.89 | 0.38 ± 0.49 | 0.39 ± 0.74 | -0.84 ± 0.36 | -0.11 ± 0.56 |
| ΔLDMC | 0.55 ± 0.29 | -0.12 ± 0.35 | 0.67 ± 0.4 | 0.95 ± 0.33 | 0.18 ± 0.34 | 1.65 ± 0.98 | 1.55 ± 0.47 | 2.53 ± 0.52 |
| ΔSLA | -0.26 ± 0.11 | 0.02 ± 0.15 | 0.02 ± 0.12 | -0.17 ± 0.11 | -0.02 ± 0.05 | -0.29 ± 0.08 | -0.04 ± 0.09 | -0.41 ± 0.08 |
| ΔC:N ratio | -0.15 ± 0.14 | 0.07 ± 0.24 | -0.17 ± 0.36 | 1.24 ± 0.9 | -2.16 ± 0.83 | -2.09 ± 0.86 | -2.31 ± 0.63 | -3.23 ± 0.64 |
| Δheight | 1.76 ± 2.12 | 1.59 ± 2.8 | 1.43 ± 1.83 | 3.03 ± 2.26 | 4.34 ± 1.16 | 7.36 ± 1.61 | 4.4 ± 0.86 | 8.21 ± 1.53 |
| CE | 1.82 ± 1.28 | 6.2 ± 1.22 | 2.95 ± 1.12 | 6.71 ± 2.17 | -0.29 ± 0.59 | 0.08 ± 0.98 | 1.81 ± 0.97 | 0.5 ± 0.49 |
| SE | 3.78 ± 1.12 | 5.34 ± 1.64 | 2.8 ± 1.2 | 4.97 ± 2.34 | 1.27 ± 0.36 | 1.5 ± 0.58 | 0.11 ± 0.34 | 0.97 ± 0.29 |
| Δyield | 55 ± 18.03 | 115.4 ± 16.8 | 57.5 ± 12.3 | 116.8 ± 23.4 | 9.9 ± 5.94 | 15.9 ± 9.47 | 19.2 ± 9.72 | 14.7 ± 5.19 |
